# Conservation of neuron-astrocyte coordinated activity among sensory processing centers of the developing brain

**DOI:** 10.1101/2024.04.15.589519

**Authors:** Vered Kellner, Patrick Parker, Xuelong Mi, Guoqiang Yu, Gesine Saher, Dwight E. Bergles

## Abstract

Afferent neurons in developing sensory organs exhibit a prolonged period of burst firing prior to the onset of sensory experience. This intrinsically generated activity propagates from the periphery through central processing centers to promote the survival and physiological maturation of neurons and refine their synaptic connectivity. Recent studies in the auditory system indicate that these bursts of action potentials also trigger metabotropic glutamate receptor-mediated calcium increases within astrocytes that are spatially and temporally correlated with neuronal events; however, it is not known if this phenomenon occurs in other sensory modalities. Here we show using *in vivo* simultaneous imaging of neuronal and astrocyte calcium activity in awake mouse pups that waves of retinal ganglion cell activity induce spatially and temporally correlated waves of astrocyte activity in the superior colliculus that depend on metabotropic glutamate receptors mGluR5 and mGluR3. Astrocyte calcium transients reliably occurred with each neuronal wave, but peaked more than one second after neuronal events. Despite differences in the temporal features of spontaneous activity in auditory and visual processing regions, individual astrocytes exhibited similar overall calcium activity patterns, providing a conserved mechanism to synchronize neuronal and astrocyte maturation within discrete sensory domains.

## Introduction

Sensory processing areas of the brain rely on a combination of genetically pre-programmed events and activity-dependent modification to assemble circuits capable of extracting features of the external world (Goodman and Shatz, 1993). The neural activity that shapes these nascent sensory circuits emerges very early in brain development (Babola et al., 2021; Bansal et al., 2000), prior to the onset of sensory experience. This intrinsically generated activity is initiated within developing sensory organs and propagates through the CNS, providing the means to establish and refine synaptic connections between peripheral and central sensory centers by inducing periods of correlated activity among neurons that will ultimately be responsible for processing similar features of sensory stimuli (Leighton and Lohmann, 2016). The mechanisms that induce this sensory-independent neuronal activity vary between sensory organs, but the resultant activity shares many similarities, consisting of discrete bursts of action potentials that repeat many times per minute over a period of several days, providing tens of thousands of discrete events with which to induce changes in gene expression and synaptic weights (Blankenship and Feller, 2010). Despite the prevalence of this early, sensory-independent activity, the mechanisms through which it shapes brain development remain poorly understood.

In the retina, burst firing of retinal ganglion cells (RGCs) is first induced by the release of acetylcholine by a subset of amacrine cells, triggering a gap-junction dependent propagating wave of correlated activity among primary sensory neurons that project to the brain (Feller et al., 1997; Meister et al., 1991; Voufo et al., 2023). Spontaneous retinal activity is highly conserved among species, having been observed in embryonic chicks (Catsicas et al., 1998; Wong et al., 1998), developing mice (Mooney et al., 1996), ferrets, cats (Meister et al., 1991), and turtles (Sernagor and Grzywacz, 1995), among others. Spontaneous neuronal activity in these nascent circuits refines projections from the retina to the lateral geniculate nucleus (LGN) (Shatz and Stryker, 1988; Sretavan et al., 1988) and to the superior colliculus (SC) (Xu et al., 2011). Moreover, the specific temporal and spatial organization of these retinal waves is required for proper axonal refinement (McLaughlin et al., 2003; Xu et al., 2015), and their directionality influences maturation of the circuits that detect motion (Ge et al., 2021), demonstrating that this early activity is important for shaping adult vision, providing a framework to understand how developing sensory organs influence maturation of neural circuits in the brain.

Although neurons provide the means for rapid information transfer, the assembly of functional circuits depends crucially on the coincident maturation of astrocytes, which secrete factors important for synapse formation and refinement (Clarke and Barres, 2013), position glutamate transporters near sites of release (Benediktsson et al., 2012; Bergles and Jahr, 1997), isolate synapses through ensheathment (Grosche et al., 1999), and regulate synaptic function through the release of transmitters (Dallérac et al., 2018). Recent studies in the auditory system of mice indicate that spontaneous bursts of action potentials initiated within the developing cochlea that are analogous to retinal waves, trigger spatially and temporally correlated activity of neurons and astrocytes within sound processing centers of the brain prior to hearing onset (Kellner et al., 2021), providing a possible means to coordinate their development. The threshold for induction of astrocyte activity is higher than for neurons, with only the strongest bursts reaching the threshold for astrocyte activation, consistent with a requirement for extrasynaptic glutamate spillover and pooling of extrasynaptic glutamate transients between nearby synapses to engage peripherally located metabotropic receptors on astrocyte processes (Bergles et al., 1999). Astrocyte increases in calcium were primarily dependent on activation of mGluR5, a metabotropic glutamate receptor that is highly expressed by astrocytes only during early brain development (Cai et al., 2000; Sun et al., 2013a). As coordinated astrocyte activity in these regions is dependent on neuronal burst firing that emerges from the cochlea, it suggests that mGluR5 may be developmentally expressed in astrocytes to enable detection of this early spontaneous activity. Although neuronal burst firing is conserved among developing sensory pathways, it is not yet known whether astrocytes are also recruited by burst firing in other developing sensory modalities. Differences in patterns of spontaneous neuronal activity between different sensory organs could lead to different spatio-temporal profiles of extrasynaptic glutamate, and therefore altered engagement of astrocytes. Moreover, astrocytes exhibit heterogeneity with regard to receptor expression and synaptic ensheathment (Khakh and Sofroniew, 2015; Ventura and Harris, 1999), which could also influence the degree of responsiveness to perisynaptic glutamate transients induced by neuronal burst firing.

To determine whether coordinated activation of neurons and astrocytes is conserved in other sensory systems, we used *in vivo* widefield and two-photon imaging of genetically encoded calcium indicators selectively expressed in astrocytes and neurons to define activity patterns in the SC, a visual processing region of the midbrain that receives direct retinal input. Simultaneous neuron and astrocyte calcium imaging in unanesthetized mouse pups revealed that burst firing of RGCs leads to co-activation of neurons and astrocytes within discrete spatial domains defined by wave propagation in the SC. Despite differences in neuronal activity patterns that emerge from visual and auditory sensory organs, individual astrocytes in both regions experienced similar overall levels of activity that was dependent on co-activation of metabotropic glutamate receptors mGluR5 and mGluR3, indicating that induction of coordinated neuron-astrocyte activity by intrinsically generated neuronal activity is conserved between brain regions and sensory modalities, providing a means to synchronize their maturation.

## Results

### Retinal waves induce coordinated increases in calcium among SC astrocytes

Groups of retinal ganglion cells (RGCs) fire bursts of action potentials that move across the retina in a wave like pattern before eyelid opening (Meister et al., 1991). This spatially coordinated activity propagates through developing visual processing circuits, eventually reaching the superior colliculus (SC) and the visual cortex (Ackman et al., 2012). We previously discovered that glutamate spillover from active synapses during spontaneous burst firing of auditory neurons coordinates activity between neurons and astrocytes in developing sound processing centers of the brain (Kellner et al., 2021). To determine if this spatial and temporal coordination of neuron and astrocyte activity is conserved among nascent sensory processing circuits, we generated mice that express the genetically encoded calcium indicator GCaMP6s in astrocytes after tamoxifen induction (*Aldh1l1-CreER;R26-lsl-GCaMP6s*; hereafter termed “*Aldh1l1;GC6s”* mice), implanted a cranial window above the midbrain, and used wide field mesoscopic epifluorescence time-lapse imaging to visualize their activity in the SC before eye opening (P6-11) (Figures 1A-1B). Widefield imaging of GCaMP6s fluorescence within the SC revealed that groups of astrocytes experienced transient increases of calcium, forming a band that propagated across the SC in a wave-like manner (Figure 1C-1D; Movie S1). We adapted the Astrocyte Quantitative Analysis (AQuA) software (Wang et al., 2019) to detect and quantify individual waves (see Methods; Figure S1). Astrocyte calcium waves occurred in the SC with an average frequency of 4.8 ± 0.59 waves/min (n = 17 mice ages P6-P11), comparable to the frequency of neuronal activity waves observed in the SC under similar recording conditions (Gribizis et al., 2019). Astrocyte calcium waves occurred in both the left and right SC with similar frequency (4.9 ± 0.7; 4.89 ± 0.6 waves/min for left and right SC, respectively; n = 17 mice ages P6-P11; Wilcoxon signed rank test, p = 0.1), but appeared to be induced independently, as there was low temporal correlation of activity between the two lobes of the SC (16.5 ± 3% of events occurred bilaterally on average; n = 9 mice ages P6-P10).

**Figure 1.**
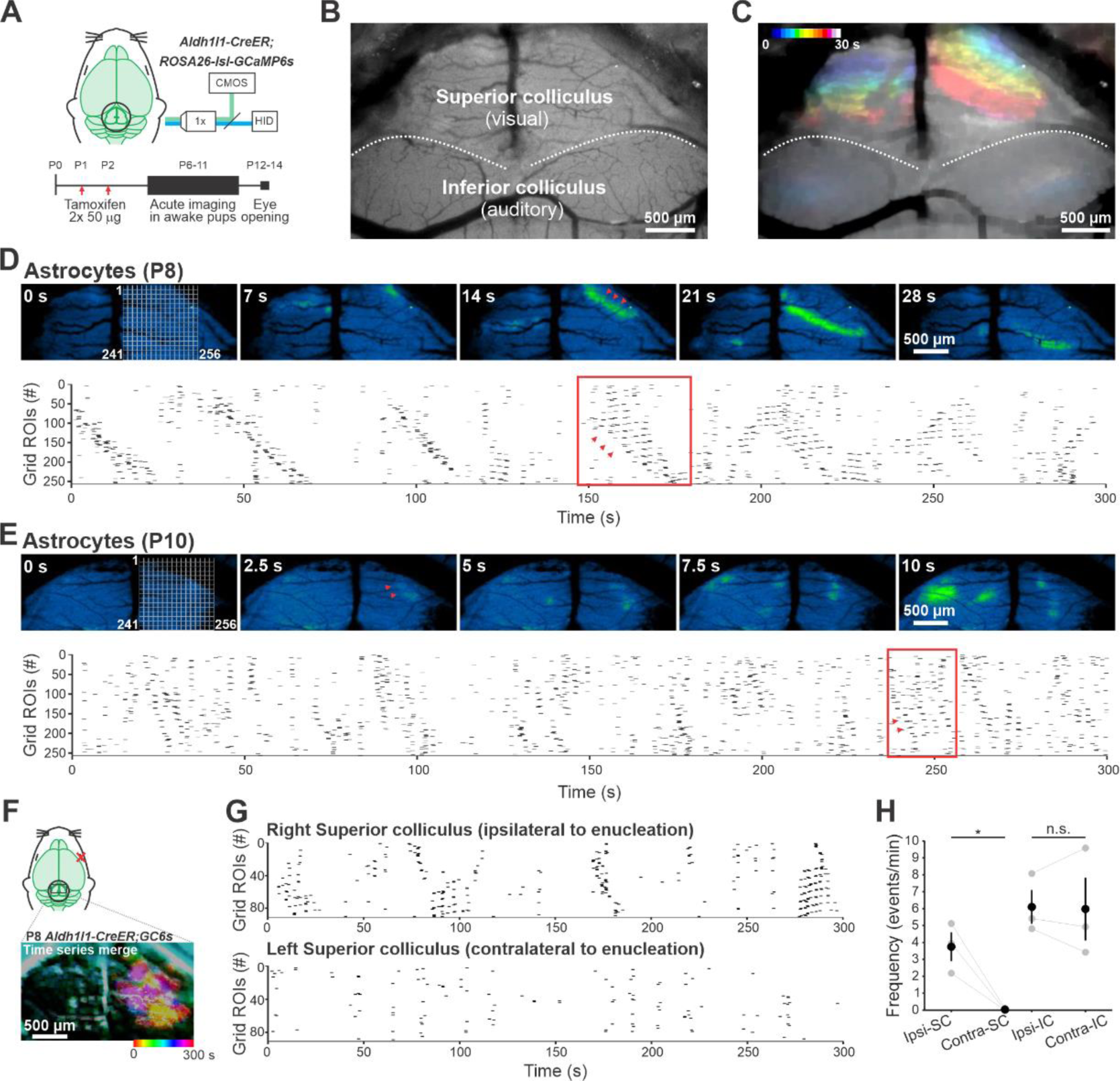
Astrocytes exhibit retinal waves before eye opening. **A.** Experimental set up and time line for experiments. **B**. View through the widefield microscope showing the superior and inferior colliculus (SC and IC, respectively) from a postnatal day 8 (P8) Aldh1l1-creER;GCaMP6s mouse. **C**. Temporal color-coding of 30 s of astrocyte activity from the same mouse shown in B. **D**. Montage of an astrocyte retinal wave occurring in the SC of the same mouse shown in B,C (Top; pseudocolored) and a raster plot of all the detected peaks from each grid shown on the right SC of the first image, over 5 minutes of imaging (Bottom). Red arrows show direction of propagation in the montage and the raster plot. **E**. Montage of several calcium events in a P10 mouse, showing that the characteristics of the waves change at this stage of development (Top; pseudocolored) and a raster plot of 5 minutes of imaging (Bottom). **F-G**. Right eye enucleation experiment shows astrocyte activity is dependent on retinal input. **F**. Illustration of the right eye enucleation (red x; Top) and a temporal color-coding of 300 s of astrocyte activity in a P7 Aldh1l1;GC6s mouse showing activity on the ipsilateral SC but not the contralateral SC (Bottom; Background subtracted for clarity). **G**. Raster plot of detected peaks from each hemisphere of the mouse shown in F. Note that remainder of events in the left SC (bottom) are none directional and associated with movements of the animal. **H**. Frequency of waves in SC and events in IC in the ipsilateral and contralateral hemispheres to the ablated eye. Gray circles represent individual animals (lines connect same animal). Colored circles represent means and scale bars represent SEMs. Paired t-test. n=3 mice (P7-P9). n.s. = not significant; *p<0.05.

The features of spontaneous neuronal activity within the SC evolve with age, reflecting functional changes in the induction and propagation of retinal waves as animals mature (Ackman et al., 2012; Gribizis et al., 2019). Neuronal waves in stage II, the ‘cholinergic stage’ (P0-P9), feature large, directional sweeps over the entire SC (Ackman et al., 2012). Astrocyte calcium waves from mice of this age exhibited similar properties, consisting of a continuous line of co-active astrocytes that propagated primarily in one direction, visible as diagonals in raster plots of cellular activity (Figure 1D; Movie S2). In contrast, neuronal waves in stage III (P10-P14), the ‘glutamatergic stage,’ are more frequent, faster, and often appear as spatially restricted wavelets, rather than long, unbroken wavefronts (Gribizis et al., 2019). At this age (P10-P14), astrocyte calcium waves were also more frequent (P6-9: 3.8 ± 0.3 waves/min, n = 13; P10-11: 7.2 ± 1.2 waves/min, n = 6; p = 0.016; Wilcoxon rank sum test) and less organized, with correlated events occurring in more restricted regions of the SC that propagated for shorter distances (Figure 1E; Movie S2). Thus, astrocyte calcium changes in the developing SC mirror changes in RGC activity prior to eye opening.

If correlated astrocyte activity in the SC is induced by glutamate release from local RGC projections, they should be dependent on retinal input. Unilateral eye enucleation prior to imaging, abolished wave like astrocyte calcium activity in the contralateral SC (P7-P9, n = 3) (Figures 1F-1H). Notably, eye enucleation did not alter propagating waves in the ipsilateral SC (P7-9 ipsilateral SC with enucleation: 3.7 ± 0.8 waves/min n = 3 compared to P6-9: 3.8 ± 0.3 waves/min, n = 13 general controls; p = 0.88, Wilcoxon rank sum test), or cochlea-driven spontaneous activity patterns in the inferior colliculus (IC) (P7-9 IC with enucleation: 5.7 ± 1.2 events/min n = 3 compared to P6-9: 7.14 ± 0.5 events/min, n = 13 general controls; p = 0.29, Wilcoxon rank sum test), suggesting that the loss of astrocyte activity was not caused by non-specific aspects of the surgery. Together, these results show that astrocytes within the developing visual midbrain exhibit correlated increases in calcium during early development that are dependent on input from the retina.

### Spatial and temporal coordination of astrocyte and neuronal activity in the developing SC

To determine the precise spatial and temporal relationship between astrocyte and neuronal activities in the SC, we generated mice that express both GCaMP6s in astrocytes and the red-shifted calcium indicator jRGECO1a in neurons (*Aldh1l1;GC6s;Thy1-jRGECO1a* mice, P6-P11) (Figures 2A-2B). Simultaneous, two color, two photon *in vivo* imaging revealed that each propagating wave of neuronal activity was followed by a wave of astrocyte activity aligned to the neuronal wave front. Astrocyte responses peaked with an average lag time of 1.4 seconds relative to the peak neuronal response (Figure 2C and 2D; Movie S3). Unbiased sampling of astrocyte fluorescence amplitude 1.5 seconds after the peak of neuronal fluorescence, revealed that the amplitudes of neuronal and astrocyte calcium changes were highly correlated (Figure 2E), indicating that unlike astrocyte activity in the IC, astrocytes in the SC were responsive to a wide range of neuronal activities, with almost every neuronal wave followed by an astrocyte wave.

**Figure 2.**
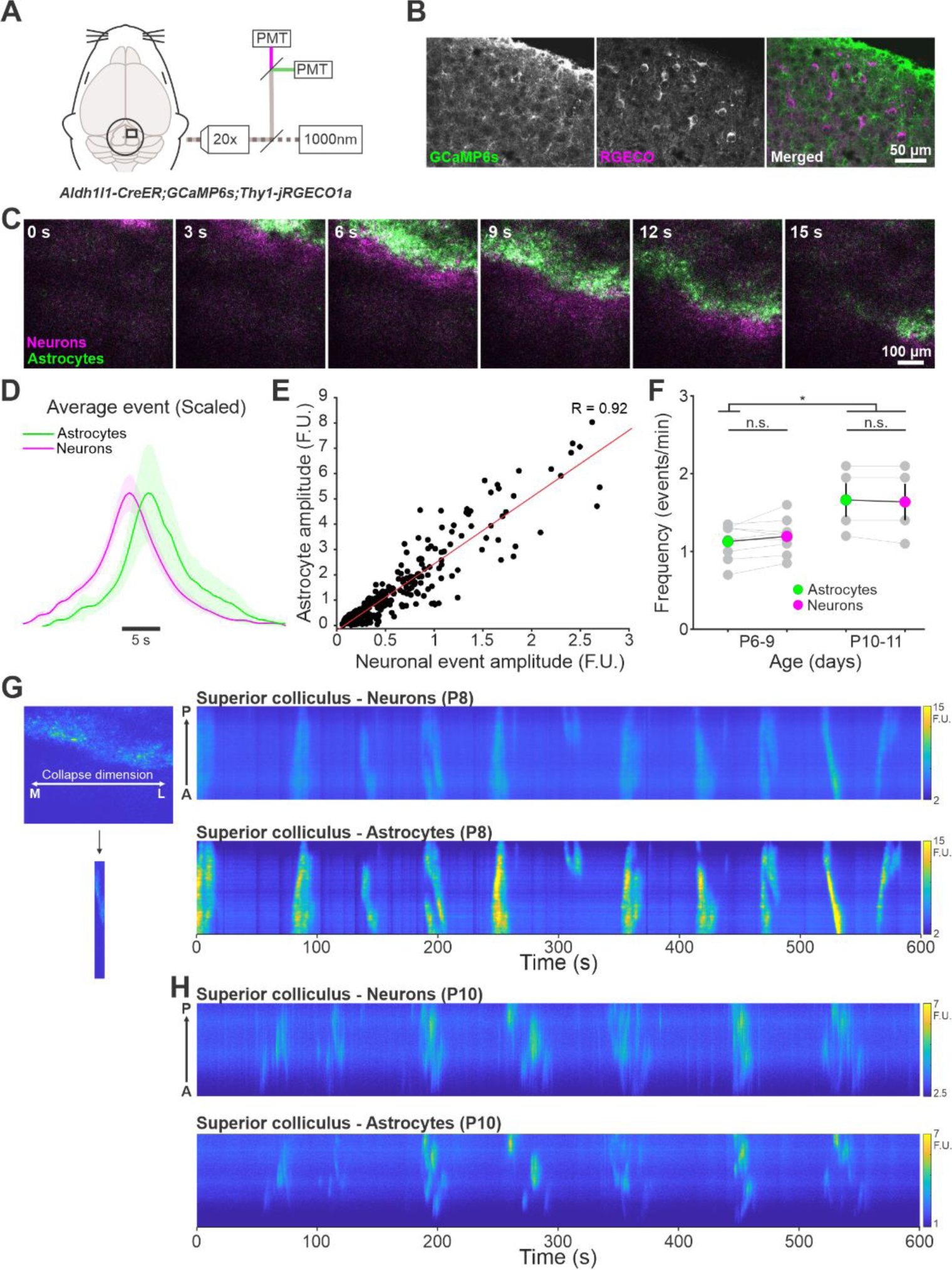
High correlation of events between astrocytes and neurons. **A**. Experimental set up for simultaneous imaging of astrocytes and neurons using a two photon microscope. **B**. Immunohistochemistry against GFP and mCherry to show the expression of GCaMP6s and jRGECO1a in the SC of a P11 Aldh1l1-creER;GC6s;Thy1-jRGECO1a mouse showing astrocytes (left), neurons (middle) and the overlap between them (right; includes DAPI stain). **C**. Montage of one retinal wave showing the increase in calcium in neurons (magenta) and astrocytes (green) from the mouse shown in B. **D**. Average waveform of all events detected using full field averaging of fluorescence from neurons (magenta) and astrocytes (green). The shift in time is also averaged over all peaks. Solid line is the mean, shaded area indicates SEM. n=13 mice (P6-P11). **E**. Astrocyte fluorescence values 1.5 s after a neuronal peak was detected (black) with regression line (red). F.U. = fluorescent units. Pearson correlation values are denoted on the figure. n=288 events from 13 mice. **F**. Retinal wave frequency in astrocytes (green) and neurons (magenta) at different postnatal ages. Gray circles represent individual animals (lines connect astrocytes and neurons in same animal). Colored circles represent means and scale bars represent SEMs. Unbalanced two-way ANOVA. n = 9, 4 mice for P6-9 and P10-11, respectively. **G**. Calcium events shown by collapsing the medial/lateral dimension and showing the events from anterior (A) to posterior (P) in neurons (top) and astrocytes (bottom) from the mouse in B,C. **H**. Same as in G from a P10 mouse showing that the nature of the waves changes during development. n.s. = not significant; *p<0.05.

Imaging activity in astrocytes and neurons at different developmental time points revealed a slight increase in the frequency of astrocyte events during stage III (P10-P11), but a persistent high correlation between astrocyte and neuron activities (Figure 2F), even at later developmental ages when neuronal events (retinal waves) were smaller and less directional (Figures 2G-2H). This tight coordination was distinct from the activity patterns in the IC induced by spontaneous activity in the cochlea. Sequential imaging of the IC and SC in the same mice (Figures 3A-3C), revealed that astrocyte activity in the IC did not occur after each neuronal event (Figure 3D, Movie S4), consistent with previous observations (Kellner et al., 2021). Moreover, both neuronal and astrocyte events in the IC were more frequent than events in the SC (for astrocytes: SC: 1.18 ± 0.09; IC: 4.3 ± 0.16 events/min, n = 5, p = 2.07*10^-4^, Paired t-test; for neurons: SC: 1.25 ± 0.1; IC: 6.9 ± 0.18 events/min, n = 5, p = 8.5*10^-6^, Paired t-test), despite the 0.5x lower frequency of astrocyte events relative to neuronal events in the IC (Figure 3E). Correspondingly, the correlation coefficients between neuronal and astrocyte signals were significantly lower in the IC than in the SC (Figures 3F). However, the delay between the peaks of neuronal and astrocyte activity was not significantly different between the two areas (Delay time: IC: 1.8 ± 0.12 s; SC: 1.5 ± 0 s, n = 5, p = 0.16, Wilcoxon rank sum test), consistent with a dependence on metabotropic glutamate receptor engagement on astrocytes. These results suggest that the different spatial and temporal features of neuronal burst firing that emerge from the developing retina and cochlea induce distinct global patterns of astrocyte activity in central visual and auditory processing centers.

**Figure 3.**
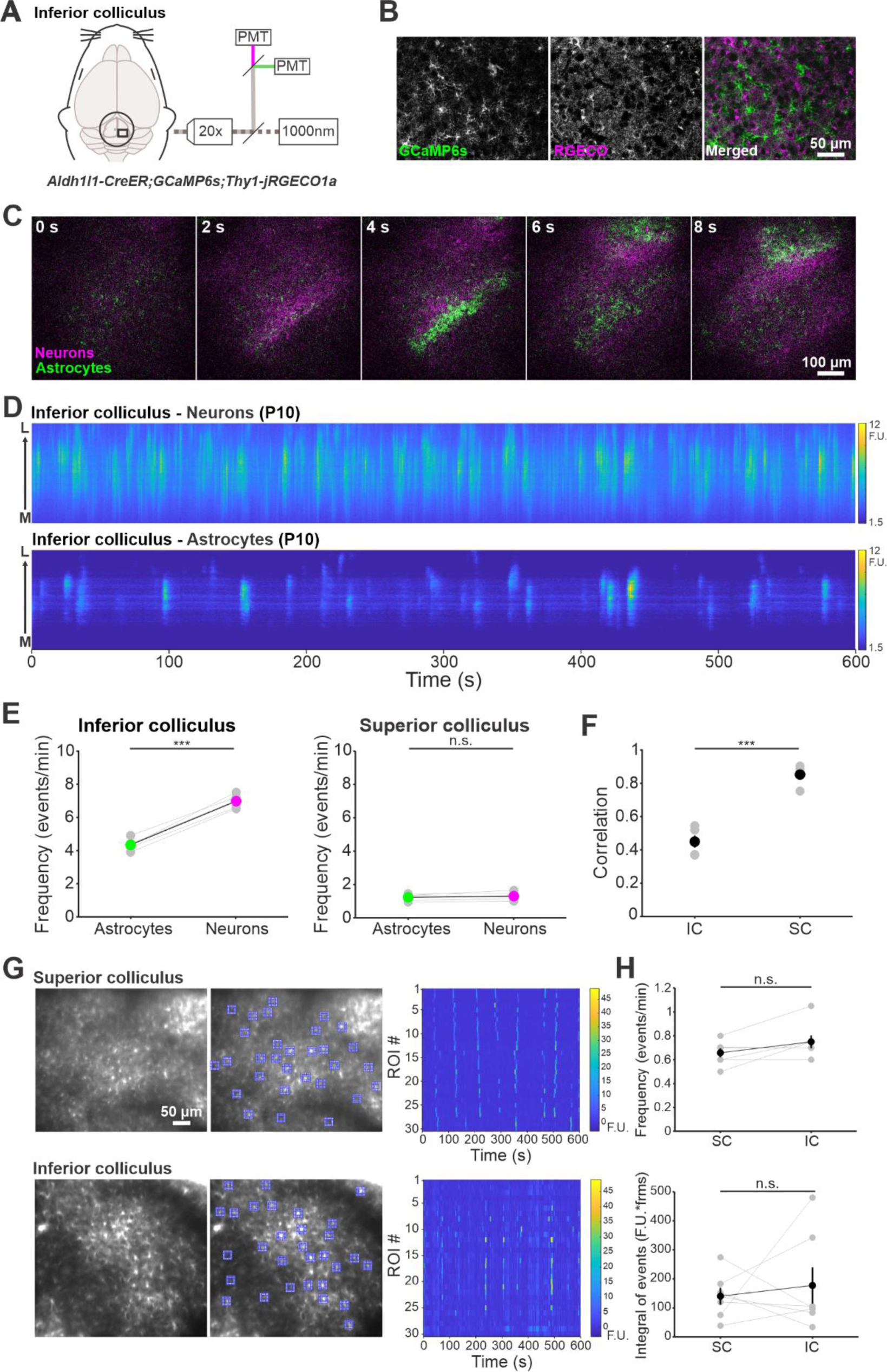
Astrocyte-neuron events in the inferior colliculus are less correlated than in the superior colliculus but single cells integrate the same amount of activity. **A**. Experimental set up for simultaneous imaging of astrocytes and neurons using a two photon microscope showing that in some animals the inferior colliculus (IC) was also imaged. **B**. Immunohistochemistry against GFP and mCherry to show the expression of GCaMP6s and jRGECO1a in the IC of a P11 Aldh1l1-creER;GCaMP6s;Thy1-jRGECO1a mouse showing astrocytes (left), neurons (middle) and the overlap between them (right; includes DAPI stain). **C**. Montage of several neuronal events (magenta) and one astrocyte event (green) from a P10 Aldh1l1-creER;GC6s;Thy1-jRGECO1a mouse. **D**. Calcium events shown by collapsing the anterior/posterior dimension and showing the events from medial (M) to lateral (L) in neurons (top) and astrocytes (bottom) from the same mouse in C. **E**. Frequency of events in astrocytes (green) and neurons (magenta) in the IC (left) and SC (right) over all postnatal ages. Note that the SC values are a subset of the values in Figure 2F combined over ages. Paired t-test; n = 5. **F**. Correlation coefficients between astrocyte and neuronal calcium traces in the IC and SC. Student’s t-test. n = 5. **G**. Average projection images over 10 minute imaging session from SC (top) and IC (bottom) of a P7 Aldh1l1;GC6s mouse (left) with 30 ROIs displayed in blue (middle) and the calcium fluorescence change over time shown for each ROI (right). F.U. = fluorescent units. **H**. Frequency of events (top) and integral of detected calcium peaks (bottom) averaged over all ROIs per mouse in the SC and IC. Lines connect averages from the same mouse. Wilcoxon sign rank test; p=0.25 and 0.94 for frequency and integral, respectively. n=7 mice. n.s.= not significant; ***p<0.001.

To determine if the macroscopic features of these spontaneous events lead to differences in the profile of calcium changes induced within individual astrocytes, we used high resolution two photon imaging to monitor somatic GCaMP6s fluorescence changes in astrocytes in these two regions (Figure 3G). Remarkably, individual astrocytes in each region displayed similar calcium activity changes, with comparable frequencies and fluorescence integrals over similar imaging times (Figure 3H). Thus, despite differences in the kinetics of intrinsically generated activity between the retina and the cochlea, astrocytes in these two developing sensory domains experience similar calcium dynamics.

### Astrocyte calcium events in the SC result from co-activation of mGluR5 and mGluR3

Our previous studies revealed that burst firing of neuronal afferents in the IC results in activation of mGluR5 and mGluR3 on astrocytes, which synergize to release calcium from intracellular stores (Kellner et al., 2021). In the IC, pharmacological inhibition of mGluR5 or selective astrocyte deletion of *Grm5* (mGluR5) was sufficient to abolish calcium transients in astrocytes initiated by cochlear activity. To determine if mGluR5 is also the dominant receptor mediating astrocyte calcium activity in the SC, we monitored astrocyte activity in mice before and after an intraperitoneal injection of an mGluR5 antagonist, MPEP (10 mg/kg). In both SC and IC, MPEP significantly reduced the astrocyte calcium events (SC: 4.7 ± 0.46 and 1.75 ± 0.45 waves/min before and after MPEP, respectively; IC: 7.6 ± 0.25 and 1.3 ± 0.6 events/min before and after MPEP, respectively; Paired t-test, p<0.001 for both). Because this pharmacological manipulation could have altered neuronal mGluR5, we then monitored astrocyte activity in mice in which mGluR5 had been conditionally deleted from astrocytes (*Aldh1l1-CreER;Grm5^fl/fl^;GC6s*; hereafter termed “mGluR5-cKO*”* mice). In mGluR5-cKO mice, the frequency of calcium waves in SC and IC were both lower than in control mice (*Aldh1l1;GC6s*); however, events in SC were only marginally significantly reduced (Figures 4A and 4B). Acute administration of the mGluR5 antagonist, MPEP to mGluR5-cKO mice, did not alter the remaining astrocyte SC activity (mGluR5-cKO: 3.3 ± 0.6 events/min; mGluR5-cKO+MPEP: 4.9 ± 1.14 waves/min, n = 2), but abolished residual astrocyte activity in the IC (mGluR5-cKO: 1.36 ± 0.6 events/min; mGluR5-cKO+MPEP: 0.02 ± 0.02 waves/min, n = 2), indicating that the residual activity in SC was not due to persistent expression of mGluR5.

**Figure 4.**
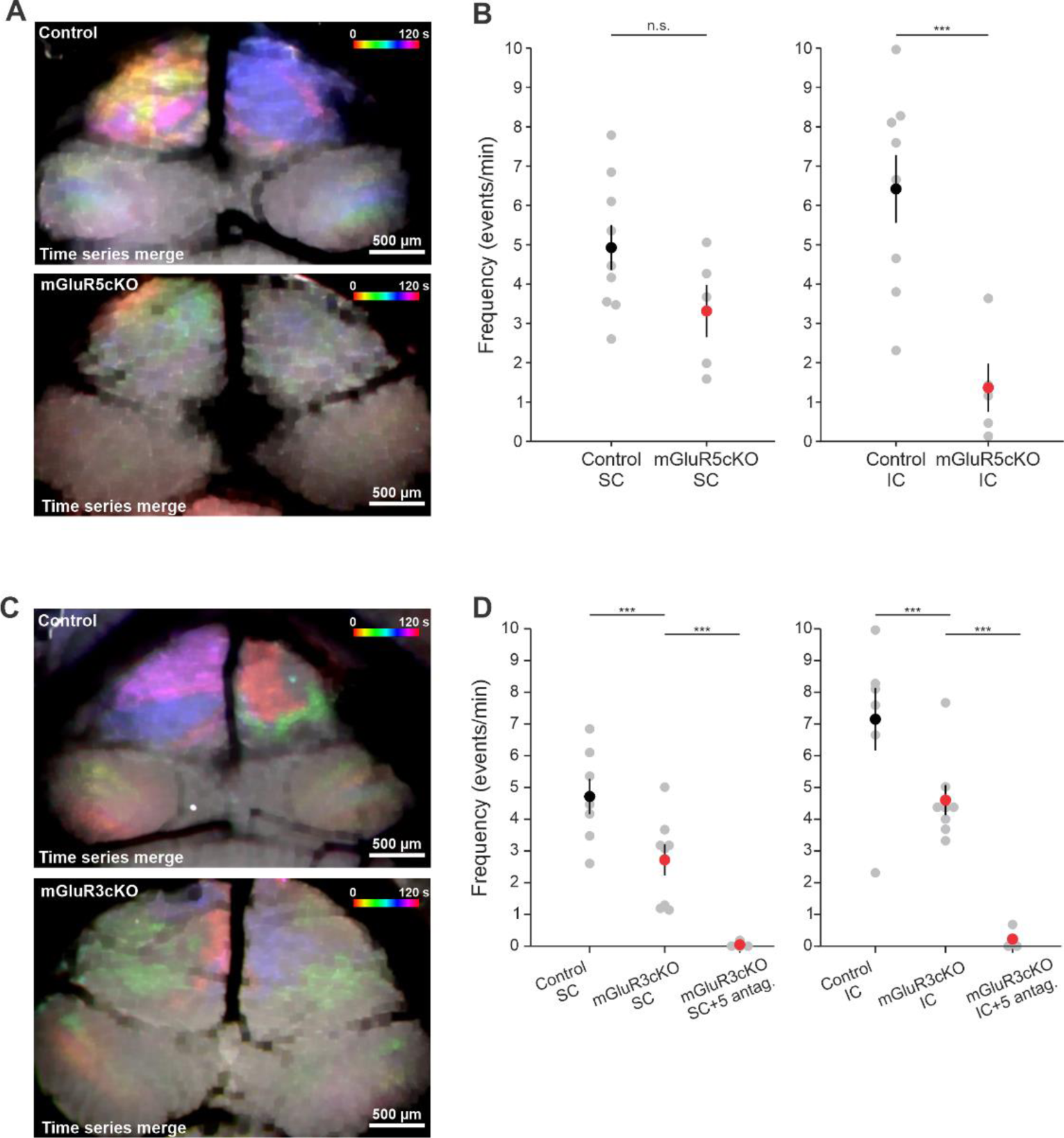
Astrocyte calcium events in the superior colliculus require both mGluR5 and mGluR3. **A**. Temporal color-coding of 120 s of astrocyte activity in a control (Aldh1l1;GC6s) P8 mouse (Top) and an mGluR5-cKO (Aldh1l1;mGluR5^fl/fl^;GC6s) P10 mouse (Bottom). **B**. Frequency of astrocyte events in SC (Left) and IC (Right) in control and in mGluR5cKO mice (same mice for SC and IC). Gray circles represent individual mice. Colored circles represent means and scale bars represent SEMs. n = 10, 5 mice for control and cKO, respectively. Student’s t-test. Note that not all cases had full recombination. **C**. Temporal color-coding of 120 s of astrocyte activity in a control (Aldh1l1;GC6s) P8 mouse (Top) and an mGluR3cKO (Aldh1l1;mGluR3^fl/fl^;GC6s) P7 mouse (Bottom). **D**. Frequency of astrocyte events in SC (Left) and IC (Right) in control, mGluR3cKO, and in the mGluR3cKO mice (same mice for SC and IC) in the presence of the mGluR5 antagonist, MPEP (10 mg/kg I.P.). Gray circles represent individual mice. Colored circles represent means and scale bars represent SEMs. n = 7, 8, 4 mice, respectively. 3-way nested ANOVA; n.s. = not significant; ***p < 0.001.

To assess the contribution of mGluR3 receptors, which contribute to intracellular calcium release in astrocytes in the IC (Kellner et al., 2021), we measured astrocyte calcium changes in mice with astrocyte-specific knockout of *Grm3* (mGluR3) (*Aldh1l1-CreER;Grm3^fl/fl^;GC6s*; hereafter termed “mGluR3-cKO*”* mice). Astrocyte calcium activity was significantly lower in both SC and IC in mGluR3-cKO mice compared to controls (Figure 4C and 4D). Administration of the mGluR2/3 antagonist, LY341495 to mGluR3-cKO mice, did not alter the frequency of astrocyte calcium waves in SC (2.7 ± 0.5 vs. 3.3 ± 0.5 waves/min, n = 2, in mGluR3-cKO and mGluR3-cKO+LY, respectively), or IC (4.6 ± 0.5 vs. 3.5 ± 0.2 events/min, n = 2, in mGluR3-cKO and mGluR3-cKO+LY, respectively), indicating efficient *Grm3* deletion. All remaining activity in both SC and IC was abolished by acute administration of MPEP in mGluR3-cKO mice (Figure 4D). Together, these results indicate that both mGluR5 and mGluR3 metabotropic glutamate receptors are used by astrocytes in these distinct sensory processing areas, albeit with slightly different impact, to detect the burst firing of sensory neurons prior to the onset of sensory experience.

## Discussion

The development and maturation of astrocytes coincides with a period of extensive synapse maturation and refinement in the CNS (Freeman, 2010). However, the physiological patterns of activity experienced by astrocytes at this critical developmental phase have not been well defined *in vivo*. Spontaneous neuronal activity that passes through nascent sensory processing centers of the brain occurs as prolonged bursts of action potentials (Mooney et al., 1996; Tritsch et al., 2010), a stimulus that has the potential to release large amounts of glutamate into the synaptic cleft during each burst. Moreover, because bursts are correlated among neurons that project to similar regions, glutamate transients from adjacent active synapses can combine, increasing the peak glutamate concentration and prolonging the decay of these burst-mediated transients near astrocyte membranes. Astrocytes are poised to sense these extrasynaptic glutamate transients through metabotropic glutamate receptors, especially mGluR5, which is highly expressed in astrocytes during development (Cai et al., 2000; Sun et al., 2013b). Here, we show using *in vivo* imaging from unanesthetized mouse pups, that burst firing of RGCs that occurs during retinal waves reliably induces calcium increases in astrocytes within the SC that closely follow the propagating wave of neuronal activity. Thus, close spatial-temporal coordination of neuron and astrocyte activity is a common feature of developing circuits involved in processing sensory information.

### Spatial and temporal coordination of neuron-astrocyte activity in the SC

If neuronal burst firing provides a conserved means to coordinate the maturation of astrocytes and neurons, one might expect that the calcium changes experienced by these cells would be similar. However, there were distinct differences in astrocyte activation between these two developing sensory circuits. In the SC, the coordination between astrocytes and neurons was remarkably consistent, with each neuronal wave being followed by an astrocyte wave (Figure 2). The mechanisms responsible for generating neuronal bursts in the retina and the cochlea provide an explanation for these different characteristics. In the cochlea, spontaneous activity is generated by the stochastic release of ATP from glia-like inner supporting cells at different locations along the sensory epithelium, initiating a cascade of events that eventually cause a group of nearby inner hair cells to depolarize and activate first order sensory neurons (spiral ganglion neurons, SGNs; Tritsch et al., 2007). Through this mechanism, correlated activity is induced in groups of neurons that that will later process similar frequencies of sound (Babola et al., 2018). Notably, this activity is spatially constrained and does not propagate along the length of the sensory epithelium. Like SGNs in the cochlea, RGC activity in the retina is similarly spatially and temporally correlated; however, RGC activity propagates as slow-moving waves, particularly during cholinergic stage II. As a result, the burst duration in RGCs is much longer than in SGNs (8 seconds vs. 5 seconds, respectively) (Sun et al., 2008; Babola et al., 2020), and the number of spikes per burst is also significantly larger (RGCs: 100 spikes/burst; SGNs: <50 spikes/burst), suggesting that the amount of glutamate spillover in the SC could be much higher than in the IC, allowing for more consistent engagement of astrocyte metabotropic receptors. However, it is possible that differences in receptor densities, their localization, and synaptic association of astrocyte membranes may also contribute to greater mGluR activation in the SC.

### Dual mGluR activation in the SC

Astrocyte calcium activity during neuronal burst firing in the IC results from engagement of both mGluR5 and mGluR3 glutamate receptors (Figure 4) (Kellner et al., 2021). Although mGluR3 is a canonical Gi coupled receptor associated with negative regulation of cAMP, prior studies in neurons have shown that this receptor can enhance IP3 dependent calcium release from intracellular stores through regulation of phospholipase C β (Di Menna et al., 2018), providing an unexpected synergy to enhance calcium signaling. As we do not yet know the precise densities and distributions of these mGluRs on astrocyte membranes relative to release sites, it is uncertain why this dual mode of signaling is used, but it is possible that this may enable translation of different patterns of activity into distinct cellular responses. Genetic and pharmacological manipulations indicate that astrocyte calcium responses in the SC are less dependent on mGluR5 than those in IC. These variations in receptor engagement could reflect differences in receptor density, localization relative to synapses, or intracellular coupling. Despite the differential engagement of these receptors in these two sensory domains, individual astrocytes exhibited similar overall calcium increases, raising the possibility that the temporal profiles of calcium may be optimized to engage specific downstream signaling mechanisms to elicit common physiological and genetic changes (Cebolla et al., 2008).

### Potential roles of developmental astrocyte calcium signaling

Previous studies in the visual system showed that altering the pattern of spontaneous activity during development disrupts eye specific segregation and refinement of connections involved in visual processing (McLaughlin et al., 2003), and that features of spontaneous activity, namely the directionality of retinal waves, instructs the organization of direction selective circuits in the brain (Ge et al., 2021). The ability of neuronal burst firing to synchronize calcium increases in astrocytes may contribute to the circuit-level changes necessary to achieve these functional abilities. Astrocytes secrete synaptogenic factors, such as thrombospondin (Christopherson et al., 2005; Eroglu et al., 2009), Hevin and SPARC (Kucukdereli et al., 2011), chordin-like 1 (Chrdl1; Blanco-Suarez et al., 2018), and Mertk (Chung et al., 2013) to promote maturation and refinement of synapses; however, it is not yet known if the expression and/or release of these factors is altered by cytosolic calcium elevation. A recent study showed that blocking neuronal input (via thalamus) to visual cortex (using *Rora-Cre;VGlut2cKO* mice) or blocking G-protein mediated intracellular astrocyte calcium signaling (using IP3R2 null mice), altered astrocyte gene expression, including expression of glipicans 4 and 6 (*Gpc4*, *Gpc6*) and *Chrdl1* in a layer-specific manner (Farhy-Tselnicker et al., 2021), supporting the hypothesis that elevation of astrocyte calcium levels may promote the maturation of excitatory synapses.

Elevation of calcium in astrocytes has also been shown to influence neurotransmitter homeostasis. Astrocyte calcium increases mediated by TRPA1 channels induce the insertion of GABA transporters (Shigetomi et al., 2011) and those mediated by mGluRs enhance glutamate transport (Devaraju et al., 2013; Mashimo et al., 2010), key features of mature tripartite synapses. Astrocytes have also been shown to release gliotransmitters like glutamate, ATP, and D-serine in a calcium dependent manner (Araque et al., 2014), which could directly modulate the frequency of action potential firing within a burst, the propagation of this activity within developing circuits, and plasticity to strengthen or weaken nascent synapses. The close spatial and temporal coordination of neuronal and astrocyte calcium transients raises the possibility that there may be underlying coincident detectors to locally restrict these changes and promote Hebbian reinforcement.

As astrocytes mature, the tips of their processes expand into fine lamella that ramify within the neuropil and form end feet on blood vessels. This elaboration of processes has been shown to depend on different neurotransmitters: glutamate acting through mGluR5 receptors (Morel et al., 2014), GABA acting through GABAB receptors (Cheng et al., 2023), and norepinephrine acting through β1-adrenergic receptors (Rosenberg et al., 2023). Thus, transient elevations in intracellular calcium may enhance process outgrowth by astrocytes locally that are then stabilized around synapses by adhesion, creating a barrier for diffusion between nearby synapses (Lawal et al., 2022) and placing transporters and signaling receptors near sites of release. Physiological and genetic analysis of the consequences of selective, acute disruption of mGluR dependent calcium signaling in astrocytes will help to define the role of this coordinated neuron-astrocyte signaling in the maturation of sensory processing circuits in the developing CNS.

## Acknowledgments

We thank Dr. Michele Pucak, Abigail Bush and Naiqing Ye for technical assistance, Terry Shelly for machining expertise, and members of the Bergles laboratory for discussions and comments on the manuscript. V.K. was supported by an NRSA grant from the NIH (F32DC017364). Funding was provided by grants from the NIH (DC008860, NS050274), a grant from the Rubenstein Fund for Hearing Research to D.E.B, and the Deutsche Forschungsgemeinschaft (SA2114/2) to G.S.

## Author contributions

Conceptualization, V.K. and D.E.B; Methodology, V.K. and P.P.; Investigation, V.K. and P.P.; Providing reagents, G.S.; Providing analysis pipeline, G.Y. and X.M.; Formal Analysis, V.K.; Writing – Original Draft, V.K. and D.E.B.; Writing – Review & Editing, V.K., P.P. and D.E.B.; All authors read and approved the final draft.

Funding Acquisition, V.K. and D.E.B.

## METHODS

### Resource availability

#### Lead contact for reagent and resource sharing

Requests for sharing resources and reagents should be directed to the Lead Contact, Dwight Bergles.

### Materials availability

This study did not generate new unique reagents.

### Data and code availability

All imaging data reported in this paper will be shared by the lead contact upon request. All original code has been deposited on github (https://github.com/Bergles-lab/Kellner-et-al-2024-source-code) and is publicly available. Any additional information required to reanalyze the data reported in this paper is available from the lead contact upon request.

### Experimental model and subject details

Male and female mouse pups between the ages of P4 to P11 were used in experiments prior to eye opening. Gender was a random variable. Mice were maintained on a mixed background. Since we used an inducible Cre-loxP system for all experiments involving astrocytes, experimental mice were injected with tamoxifen (see Tamoxifen injections section below). Mice were housed on a 12-hour light/dark cycle and were provided food ad libitum. All experiments and procedures were approved by the Johns Hopkins Institutional Care and Use Committee.

### Transgenic Animal Models

Generation and genotyping of the following mouse lines: *Aldh1l1-CreER* (Winchenbach et al., 2016), *ROSA26-lsl-GCaMP6s* (Kim et al., 2016), *Thy1-jRGECO1a-WPRE* (GP8.62; Dana et al., 2016), *mGluR5^fl/fl^* mice (Xu et al., 2009) and *Grm3^fl/fl^*mice (Kellner et al., 2021) have been previously described.

### Method Details

#### Tamoxifen Injections

The tamoxifen solution for injections (1 mg/mL) was freshly prepared by sonicating tamoxifen (T5648, Sigma-Aldrich) in sunflower seed oil (S5007, Sigma-Aldrich) at room temperature for 20-30 min (with intermittent 20 s vortexing every 10 min). This solution was stored at 4°C for 5-7 days away from light. In *Aldh1l1-CreER;mGluR5^fl/fl^;GCaMP6s* and *Aldh1l1-CreER;mGluR3^fl/fl^;GCaMP6s* mice (Z)-4-Hydroxytamoxifen (4-OHT; H7904; Sigma-Aldrich) was used. Due to the need for triple transgenics, littermates were not used as controls but a separate cohort of *Aldh1l1-CreER;ROSA26-lsl-GCaMP6s* mice were injected with 4-OHT to serve as controls for these mice (see Fig 4). To prepare 4-OHT, powder was dissolved in 100% EtOH to achieve a final concentration of 20 mg/ml. Stock aliquots containing 50 µl were made and stored at - 80° until needed. When needed, 250 µl of sunflower seed oil was added to the aliquot, and sonicated for 20-30 min (with intermittent 20 s vortexing every 10 min). The solution was then spun using a speed vacuum to ensure evaporation of the EtOH and used within 24hrs. Mice were injected intraperitoneally (i.p.) with 50 µl of tamoxifen (50 µg per animal) or 4-OHT (200 µg per animal) between P1-P3 (unless otherwise mentioned in the text), once a day for two consecutive days, with each injection a minimum of 20 hrs apart.

#### Installation of cranial windows

The surgery was described previously in Babola *et al*., 2018 and Kellner et al., 2021. Briefly, animals were anesthetized with isoflurane (3-4% for induction, until mice were unresponsive to toe-pinch and 1-2% during the procedure). Following a local injection of lidocaine, a midline incision was made and the scalp was removed. A head bar was secured to the head using super glue (Krazy Glue). Connective tissue and muscles were either cut or pushed away above the interparietal bone. HEPES-buffered artificial cerebrospinal fluid (HEPES-aCSF) or cold PBS was applied to the exposed bone and replenished throughout the surgery. The interparietal bone was removed to expose the midbrain. The dura mater was removed, exposing the colliculi and extensive vasculature. A 5mm coverslip (CS-5R; Warner Instruments) was placed over the craniotomy, the surrounding bone was dried and super glue sealed the coverslip to the skull. Post-surgery, 0.9% NaCl solution was injected subcutaneously. The isoflurane was discontinued and the pups were placed under a warming lamp, for a minimum of 1 hour prior to imaging. After imaging was completed, all animals were euthanized and in some cases the brain was harvested.

#### *In vivo* calcium imaging

After recovering from anesthesia for 1-4 hours, pups were injected subcutaneously with lidocaine and mounted onto a metal bar to ensure head-fixation. They were maintained at 37°C by using a heating pad during imaging (TC-1000; CWE). The pups were mostly calm during imaging sessions but occasional movements occurred. These movements were detected and removed during image processing (see below). Widefield epifluorescence imaging was performed using a Hamamatsu ORCA-Flash4.0 LT digital CMOS camera coupled to a Zeiss Axio Zoom V16 stereo zoom microscope. For midbrain imaging, a 4x4mm field of view was illuminated continuously with a metal halide lamp (Zeiss Illuminator HXP 200C) and visualized through a 1X PlanNeoFluar Z 1.0x objective at 17x zoom. Image resolution was 512x512 pixels (16-bit pixel depth) after 2x2 binning to increase sensitivity. Imaging sessions consisted of uninterrupted acquisition over 5-10 minutes with an acquisition frame rate of 10 Hz, unless otherwise noted. For dual-color two photon imaging of neurons and astrocytes with single-cell resolution, a two photon microscope (Zeiss LSM 710) was used. Two photon excitation was achieved using a Ti:sapphire laser (Chameleon Ultra II; Coherent) tuned to 1000 nm. A 607x607 µm field of view was visualized using a 20x Zeiss 1.0 NA objective. Images were collected at a resolution of 512x512 pixels (8-bit pixel depth) at 2 Hz using a galvo-galvo scanner.

#### Drug delivery

For *in vivo* drug delivery, the animals were imaged as described above and then injected intraperitonially with either MPEP hydrochloride (10 mg/kg; Tocris; dissolved in ddH2O), an mGluR5 antagonist, or LY341495 hydrochloride (3 mg/kg; Tocris; dissolved in ddH2O and diluted in 0.9% NaCl), an mGluR2/3 antagonist. The animals were imaged immediately after injection (in the case of MPEP) or within 30-60 min after injection (in the case of LY341495).

#### Image processing

Widefield videos were visually inspected for clarity of the window. If the window was obstructed by blood, the movie would not be processed further. In some cases, either the right or left SC was clear and the movie was processed only for that side. For presentation of movies and in some cases (like in Figure 1F), the bleach corrected images were run through an algorithm for structured low-rank matrix factorization (Haeffele et al., 2014), this is useful in cases when the signal to noise is quite low. The ‘clean’ movie was then used for the raster plot shown in Figure 1F.

***Image Montage*** (as in Figure1): Widefield raw images were imported into Fiji as Hyperstacks, rotated, pseudocolored blue-green and adjusted for brightness and contrast. SCs were cropped and a small portion of the movie with a distinct wave was manually selected. Montage was made using the ‘Make Montage’ function.

***Raster plots*** (as in Figure1): Widefield raw images were imported into MATLAB (MATHWORKS) and corrected for photobleaching by subtracting a fitted exponent from the signal. The images were manually cropped to include either the left or right SC. Image intensities were normalized as ΔF/Fo values, where ΔF = F-Fo and Fo was defined as the median over all frames. The cropped image was then divided into 10x10 pixel squares (ROIs) and the fluorescence was averaged for each square. The calcium trace was then smoothed and filtered by regressing the baseline over multiple shifted windows using a spline approximation (‘msbackadj’ function in MATLAB, window and step size - 20 frames). Peaks in the signals were detected in MATLAB using the built-in peak detection function (‘findpeaks’) using the median + 3xMAD (median absolute deviation) as a threshold, minimum peak width of 15 frames and minimum peak prominence as half the threshold. Movement was detected using efficient subpixel image registration (Guizar-Sicairos et al., 2008) and the signal was zeroed during the detected movement events. However, this method did not detect small micromovements of the brain and so we removed detected peaks that appeared in exactly the same frame for more than 3 ROIs. The raster plot was then visualized using the function ‘plotSpikeRaster’ (Jeffrey Chiou, 2023).

***Temporal color-coded image*** (as in Figures 1C,1F,4): Widefield raw images were imported into Fiji as Hyperstacks, rotated and adjusted for brightness and contrast. The field of view was cropped and a small portion of the movie was manually selected. Background was subtracted using the ‘Subtract Background’ function with a rolling ball radius of 5 pixels and the Light background and Create background boxes checked. The ‘Temporal-Color Code’ function was used on the stack with the colormap ‘Spectrum’. Brightness and contrast were adjusted to emphasis the colors.

***Widefield tracking of calcium waves***: We applied a tracking-by-detection framework for extracting fast-moving waves from superior colliculus data. The candidate regions of waves on each frame were detected first according to the local intensity contrast, then they were associated together into different waves based on their position information.

Before candidate region detection, we used simplified Anscombe transformation 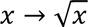 to transform Poisson noise into approximate Gaussian noise and then smoothed the data spatially with a Gaussian filter to improve the signal-to-noise ratio. Considering the size of expected waves, smoothing parameter was set to 5. Then, we adopted piecewise-linear function to model the baseline changing *F*_0_. The function was obtained by linking minimum points of equal-length segments (the length can be adjusted according to data) on the moving averaged pixel’s curve (the window for moving average was 10). Dynamic changing of intensity *dF* could be obtained.

Based on *dF*, a SynQuant-like (Wang et al., 2020) method was used to detect candidate regions of waves on each frame. In the method, a multiple-threshold strategy combined with prior knowledge (region size was limited from 100 pixels to 5000 pixels and the circularity was limited larger than 0.05) was used to pick candidate regions, whose significances were evaluated later by order-statistic test based on local intensity contrast. Among the overlapped regions selected under different thresholds, only the most significant one would be kept. A significance threshold would be further used to remove false positives, where the criteria were adjusted up to the data quality. Given candidate regions on each frame, we applied CINDA (Wang et al., 2022), an algorithm of multi-object tracking for linking regions into waves. It formulates the tracking as a minimum-cost circulation problem, as shown in Supp. Fig. 1A, where each node represents one candidate region. In the obtained flow circulation, each trajectory represents one superior colliculus wave. In our design, we set the nodes to hold the same constant reward *C*, and the edges starting from source or linking to source were set with cost 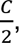 where *C* was assigned according to data quality. Specifically, the trajectory containing only one node has no contribution to the final cost, which is meaningless and would not be considered. Thus, these settings help limit the number of waves. Besides, the cost of edge linking to region A and region B in adjacent frames was set as − log(*IoU*(*A*, *B*)), where *IoU* is intersection over union, a similarity measure. It helps constrain the continuity of a wave. The linkage between nonadjacent frames with extra penalty was allowed to prevent the case that some regions were not detected during wave propagation.

Based on the prior knowledge of wave duration, a filter could be further applied in order to remove possible artifacts. According to the temporal resolution, we set the duration limitation of at least 20 frames. The movies were then inspected visually and unwanted event detections were removed (see Supp. Fig. 1B). This data was used to determine the frequency of waves as shown in Figures 1H and 4.

***Two-photon imaging***: For two-photon imaging of astrocyte and neurons simultaneously, images were imported into Fiji for presentation. The red and green channels were split and were used for further processing in MATLAB. For spatial representation of fluorescence changes, as in Figures 2G-H, the raw movie was used and one dimension was averaged to show the change in fluorescence as in a line scan over time. Colormap axes were adjusted to match between neurons and astrocytes when possible or individually when not possible. For determination of peaks, the entire image was averaged for astrocytes or neurons separately and the calcium trace was filtered by regressing the baseline over multiple shifted windows using a spline approximation (‘msbackadj’ function in MATLAB, window and step size - 45 frames). For analysis of SC data, peaks in the signals were detected in MATLAB using the built-in peak detection function (‘findpeaks’) using the median + 1xMAD/1.5 (median absolute deviation) as a threshold (thresholds were manually inspected to ensure they detected user-identified peaks), minimum peak width of 5 frames, minimum peak distance of 20 frames and minimum peak prominence as half the threshold. IC data required different parameters as the nature of the signal is different. The threshold was the median of the signal, minimum peak width of 1 frames, minimum peak distance of 5 frames (only for astrocytes, not for neurons) and minimum peak prominence as half the threshold. To calculate the lag between neuronal and astrocytic events, a cross correlation on the entire signal was performed and the lag calculated and averaged over all animals.

In some cases, as in Figure 3G-H, we manually selected ROIs within the two-photon data. The user placed 30 20x20 pixel ROIs on cell bodies. The fluorescence from these ROIs was averaged and filtered as described above for the entire field of view. For analysis of astrocytes, peaks in the signals were detected in MATLAB using the built-in peak detection function (‘findpeaks’) using the median + 3xMAD (median absolute deviation) as a threshold (thresholds were manually inspected to ensure they detected user-identified peaks), minimum peak width of 3 frames and minimum peak prominence as the threshold. For neurons, SC and IC required different thresholds - median + 3xMAD for SC and median + 1xMAD for IC, with no restrictions on minimum peak width.

***Immunohistochemistry***: Mice were deeply anesthetized with isoflurane and perfused with freshly prepared 4% paraformaldehyde (PFA) in 0.1M phosphate buffered saline (PBS). Brains were isolated and post-fixed in 4% PFA at least 4 hours and often overnight at 4°C and stored at 4°C in PBS or 30% Sucrose until processing. Brains were frozen in Tissue-Tek O.C.T. (Sakura) and cut on a cryostat microtome (Microm HM550; Thermo Fisher Scientific) into 35-50 µm sagittal or coronal sections. Before immunostaining, free-floating sections were rinsed 3x for 5 min each in PBS, then incubated for 1 hr at room temperature (RT) in a blocking buffer (0.3% Triton X-100 and 5% normal donkey serum in PBS). For immunolabeling, sections were incubated with primary antibodies diluted in blocking buffer overnight at 4°C on an orbital shaker. After rinsing at RT with PBS 3x for 10 min each, sections were incubated on an orbital shaker with fluorescent dye-conjugated secondary antibodies diluted in blocking buffer for 2-3 hr at RT. The sections were then rinsed 3x in PBS for 5 min each (during the second rinse, DAPI was added to the PBS at 1:2500 for nuclear staining) and then mounted on charged glass-slides and cover-slipped in Aqua Poly/Mount (Polysciences, Inc.). Primary antibodies used: chicken anti-GFP (1:4000, Aves Lab, Inc.), goat anti-mCherry (1:5000, SicGen). Secondary antibodies (raised in donkey): Alexa Fluor 488-, Alexa Fluor 546-, conjugated secondary antibodies to chicken or goat (1:1000, Thermo Fisher Scientific and Jackson ImmunoResearch). Images were acquired using a Zeiss LSM 800 confocal microscope with a 20x (Plan-Apochromat, Zeiss) air objective, and the pinhole set to 1 airy unit. Images were then opened with Fiji and brightness and contrast levels were adjusted. The images were pseudocolored when two or more channels were merged.

### Quantification and statistical analyses

Statistics were performed in MATLAB (Mathworks) or in Excel (Microsoft). All statistical details, including the exact value of n, what n represents, and which statistical test was performed, can be found in the figure legends or in the results section. Unless otherwise noted, data are presented as mean ± standard error of the mean. All datasets were tested for Gaussian normality using the Lilliefors test (this test was chosen because it does not require specification of the null distribution). If datasets were normal, two-tailed paired or unpaired t-tests were used. Before unpaired t-tests, the data was tested for equality of variance using the F-test. If normality was rejected, the nonparametric Wilcoxon test was used. For single comparisons, significance was defined as p < 0.05. When multiple comparisons were made, the Bonferroni correction was used to adjust p-values accordingly to lower the probability of type I errors.

## Supplemental items legend

**Movie S1: Patterned astrocyte calcium activity in the developing superior colliculus, related to Figure 1**. Astrocyte calcium waves in the superior colliculus of an awake, unanesthetized mouse (*Aldh1l1-CreER;ROSA26-lsl-GCaMP6s*) before hearing onset (P8). The movie is unprocessed and unfiltered. Images were acquired at 10 Hz; playback is 2x real time.

**Movie S2: Comparison between stage 2 and stage 3 waves in the superior colliculus, related to Figure 1**. Astrocyte calcium waves in the left superior colliculus of a P8 mouse (representing stage 2) and a P10 mouse (representing stage 3), showing the spatial and temporal differences between the stages. Images were pseudo-colored blue-green to enhance the contrast but otherwise were unprocessed.

**Movie S3: Dual-color imaging of astrocytes and neurons in the superior colliculus, related to Figure 2**. Simultaneous high resolution two photon imaging (1000 nm wavelength) of neurons (jRGECO1a; *magenta*) and astrocytes (GCaMP6s; *green*) in the right SC (102 µm depth) of an awake, unanesthetized mouse (*Thy1-jRGECO1a;Aldh1l1-CreER;R26-lsl-GCaMP6s*) before hearing onset (P8). Images were acquired at 2 Hz; playback 5x real time.

**Movie S4: Dual-color imaging of astrocytes and neurons in the inferior colliculus, related to Figure 3**. Simultaneous high resolution two photon imaging (1000 nm wavelength) of neurons (jRGECO1a; *magenta*) and astrocytes (GCaMP6s; *green*) in the right IC (180 µm depth) of an awake, unanesthetized mouse (*Thy1-jRGECO1a;Aldh1l1-CreER;R26-lsl-GCaMP6s*) before hearing onset (P10). Images were acquired at 2 Hz; playback 5x real time.

## References

Ackman, J.B., Burbridge, T.J., and Crair, M.C. (2012). Retinal waves coordinate patterned activity throughout the developing visual system. Nat. 2012 4907419 490, 219–225.

Araque, A., Carmignoto, G., Haydon, P.G., Oliet, S.H.R., Robitaille, R., and Volterra, A. (2014). Gliotransmitters Travel in Time and Space. Neuron 81, 728–739.

Babola, T.A., Li, S., Gribizis, A., Lee, B.J., Issa, J.B., Wang, H.C., Crair, M.C., and Bergles, D.E. (2018). Homeostatic Control of Spontaneous Activity in the Developing Auditory System. Neuron 99, 511–524.e5.

Babola, T.A., Li, S., Wang, Z., Kersbergen, C.J., Elgoyhen, A.B., Coate, T.M., and Bergles, D.E. (2021). Purinergic Signaling Controls Spontaneous Activity in the Auditory System throughout Early Development. J. Neurosci. 41, 594–612.

Bansal, A., Singer, J.H., Hwang, B.J., Xu, W., Beaudet, A., and Feller, M.B. (2000). Mice Lacking Specific Nicotinic Acetylcholine Receptor Subunits Exhibit Dramatically Altered Spontaneous Activity Patterns and Reveal a Limited Role for Retinal Waves in Forming ON and OFF Circuits in the Inner Retina. J. Neurosci. 20, 7672–7681.

Benediktsson, A.M., Marrs, G.S., Tu, J.C., Worley, P.F., Rothstein, J.D., Bergles, D.E., and Dailey, M.E. (2012). Neuronal activity regulates glutamate transporter dynamics in developing astrocytes. Glia 60, 175–188.

Bergles, D.E., and Jahr, C.E. (1997). Synaptic activation of glutamate transporters in hippocampal astrocytes. Neuron 19, 1297–1308.

Bergles, D.E., Diamond, J.S., and Jahr, C.E. (1999). Clearance of glutamate inside the synapse and beyond. Curr. Opin. Neurobiol. 9, 293–298.

Blanco-Suarez, E., Liu, T.-F.F., Kopelevich, A., and Allen, N.J. (2018). Astrocyte-Secreted Chordin-like 1 Drives Synapse Maturation and Limits Plasticity by Increasing Synaptic GluA2 AMPA Receptors. Neuron 100, 1116–1132.e13.

Blankenship, A.G., and Feller, M.B. (2010). Mechanisms underlying spontaneous patterned activity in developing neural circuits. Nat. Rev. Neurosci. 11, 18–29.

Cai, Z., Schools, G.P., and Kimelberg, H.K. (2000). Metabotropic glutamate receptors in acutely isolated hippocampal astrocytes: Developmental changes of mGluR5 mRNA and functional expression. 29, 70–80.

Catsicas, M., Bonness, V., Becker, D., and Mobbs, P. (1998). Spontaneous Ca2+ transients and their transmission in the developing chick retina. Curr. Biol. 8, 283–288.

Cebolla, B., Fernández-Pérez, A., Perea, G., Araque, A., and Vallejo, M. (2008). DREAM Mediates cAMP-Dependent, Ca2+-Induced Stimulation of GFAP Gene Expression and Regulates Cortical Astrogliogenesis. J. Neurosci. 28, 6703–6713.

Cheng, Y.-T., Luna-Figueroa, E., Woo, J., Chen, H.-C., Lee, Z.-F., Harmanci, A.S., and Deneen, B. (2023). Inhibitory input directs astrocyte morphogenesis through glial GABABR. Nature.

Christopherson, K.S., Ullian, E.M., Stokes, C.C.A., Mullowney, C.E., Hell, J.W., Agah, A., Lawler, J., Mosher, D.F., Bornstein, P., and Barres, B.A. (2005). Thrombospondins are astrocyte-secreted proteins that promote CNS synaptogenesis. Cell 120, 421–433.

Chung, W.S., Clarke, L.E., Wang, G.X., Stafford, B.K., Sher, A., Chakraborty, C., Joung, J., Foo, L.C., Thompson, A., Chen, C., et al. (2013). Astrocytes mediate synapse elimination through MEGF10 and MERTK pathways. Nat. 2013 5047480 504, 394–400.

Clarke, L.E., and Barres, B.A. (2013). Emerging roles of astrocytes in neural circuit development. Nat. Rev. Neurosci. 2013 145 14, 311–321.

Dallérac, G., Zapata, J., and Rouach, N. (2018). Versatile control of synaptic circuits by astrocytes: where, when and how? Nat. Rev. Neurosci. 1.

Dana, H., Mohar, B., Sun, Y., Narayan, S., Gordus, A., Hasseman, J.P., Tsegaye, G., Holt, G.T., Hu, A., Walpita, D., et al. (2016). Sensitive red protein calcium indicators for imaging neural activity. Elife 5.

Devaraju, P., Sun, M.-Y., Myers, T.L., Lauderdale, K., and Fiacco, T.A. (2013). Astrocytic group I mGluR-dependent potentiation of astrocytic glutamate and potassium uptake. J. Neurophysiol. 109, 2404–2414.

Eroglu, Ç., Allen, N.J., Susman, M.W., O’Rourke, N.A., Park, C.Y., Özkan, E., Chakraborty, C., Mulinyawe, S.B., Annis, D.S., Huberman, A.D., et al. (2009). Gabapentin Receptor α2δ-1 Is a Neuronal Thrombospondin Receptor Responsible for Excitatory CNS Synaptogenesis. Cell 139, 380–392.

Feller, M.B., Butts, D.A., Aaron, H.L., Rokhsar, D.S., and Shatz, C.J. (1997). Dynamic Processes Shape Spatiotemporal Properties of Retinal Waves. Neuron 19, 293–306.

Freeman, M.R. (2010). Specification and morphogenesis of astrocytes. Science (80-.). 330, 774–778.

Ge, X., Zhang, K., Gribizis, A., Hamodi, A.S., Sabino, A.M., and Crair, M.C. (2021). Retinal waves prime visual motion detection by simulating future optic flow. Science (80-.). 373, eabd0830.

Goodman, C.S., and Shatz, C.J. (1993). Developmental mechanisms that generate precise patterns of neuronal connectivity. Cell 72 Suppl, 77–98.

Gribizis, A., Ge, X., Daigle, T.L., Ackman, J.B., Zeng, H., Lee, D., and Crair, M.C. (2019). Visual Cortex Gains Independence from Peripheral Drive before Eye Opening. Neuron 104, 711–723.e3.

Grosche, J., Matyash, V., Möller, T., Verkhratsky, A., Reichenbach, A., and Kettenmann, H. (1999). Microdomains for neuron-glia interaction: parallel fiber signaling to Bergmann glial cells. Nat. Neurosci. 2, 139– 143.

Guizar-Sicairos, M., Thurman, S.T., and Fienup, J.R. (2008). Efficient subpixel image registration algorithms. Opt. Lett. 33, 156.

Haeffele, B.D., Young, E.D., and Vidal, R. (2014). Structured Low-Rank Matrix Factorization: Optimality, Algorithm, and Applications to Image Processing. (PMLR), pp. 2007–2015.

Kellner, V., Kersbergen, C.J., Li, S., Babola, T.A., Saher, G., and Bergles, D.E. (2021). Dual metabotropic glutamate receptor signaling enables coordination of astrocyte and neuron activity in developing sensory domains. Neuron 109, 2545–2555.e7.

Khakh, B.S., and Sofroniew, M. V (2015). Diversity of astrocyte functions and phenotypes in neural circuits. Nat. Neurosci. 18, 942–952.

Kim, Y.S., Anderson, M., Park, K., Zheng, Q., Agarwal, A., Gong, C., Saijilafu, Young, L.A., He, S., LaVinka, P.C., et al. (2016). Coupled Activation of Primary Sensory Neurons Contributes to Chronic Pain. Neuron 91, 1085–1096.

Kucukdereli, H., Allen, N.J., Lee, A.T., Feng, A., Ozlu, M.I., Conatser, L.M., Chakraborty, C., Workman, G., Weaver, M., Sage, E.H., et al. (2011). Control of excitatory CNS synaptogenesis by astrocyte-secreted proteins Hevin and SPARC. Proc. Natl. Acad. Sci. U. S. A. 108, E440–9.

Lawal, O., Ulloa Severino, F.P., and Eroglu, C. (2022). The role of astrocyte structural plasticity in regulating neural circuit function and behavior. Glia.

Leighton, A.H., and Lohmann, C. (2016). The wiring of developing sensory circuits—From patterned spontaneous activity to synaptic plasticity mechanisms. Front. Neural Circuits 10, 214646.

Madisen, L., Garner, A.R., Shimaoka, D., Chuong, A.S., Klapoetke, N.C., Li, L., van der Bourg, A., Niino, Y., Egolf, L., Monetti, C., et al. (2015). Transgenic mice for intersectional targeting of neural sensors and effectors with high specificity and performance. Neuron 85, 942–958.

Mashimo, M., Okubo, Y., Yamazawa, T., Yamasaki, M., Watanabe, M., Murayama, T., and Iino, M. (2010). Inositol 1,4,5-trisphosphate signaling maintains the activity of glutamate uptake in Bergmann glia. Eur. J. Neurosci. 32, 1668–1677.

McLaughlin, T., Torborg, C.L., Feller, M.B., and O’Leary, D.D.M. (2003). Retinotopic map refinement requires spontaneous retinal waves during a brief critical period of development. Neuron 40, 1147–1160.

Meister, M., Wong, R.O.L., Baylor, D.A., and Shatz, C.J. (1991). Synchronous Bursts of Action Potentials in Ganglion Cells of the Developing Mammalian Retina. Science (80-.). 252, 939–943.

Di Menna, L., Joffe, M.E., Iacovelli, L., Orlando, R., Lindsley, C.W., Mairesse, J., Gressèns, P., Cannella, M., Caraci, F., Copani, A., et al. (2018). Functional partnership between mGlu3 and mGlu5 metabotropic glutamate receptors in the central nervous system. Neuropharmacology 128, 301–313.

Mooney, R., Penn, A.A., Gallego, R., and Shatz, C.J. (1996). Thalamic Relay of Spontaneous Retinal Activity Prior to Vision. Neuron 17, 863–874.

Morel, L., Higashimori, H., Tolman, M., and Yang, Y. (2014). VGluT1+ Neuronal Glutamatergic signaling regulates postnatal developmental maturation of cortical protoplasmic astroglia. J. Neurosci. 34, 10950–10962.

Rosenberg, M.F., Godoy, M.I., Wade, S.D., Paredes, M.F., Zhang, Y., and Molofsky, A. V. (2023). β-adrenergic signaling promotes morphological maturation of astrocytes in female mice. J. Neurosci. JN-RM-0357–23.

Sernagor, E., and Grzywacz, N.M. (1995). Emergence of complex receptive field properties of ganglion cells in the developing turtle retina. 10.1152/Jn.1995.73.4.1355 73, 1355–1364.

Shatz, C.J., and Stryker, M.P. (1988). Prenatal Tetrodotoxin Infusion Blocks Segregation of Retinogeniculate Afferents. Science (80-.). 242, 87–89.

Shigetomi, E., Tong, X., Kwan, K.Y., Corey, D.P., and Khakh, B.S. (2011). TRPA1 channels regulate astrocyte resting calcium and inhibitory synapse efficacy through GAT-3. Nat. Neurosci. 15, 70–80.

Sretavan, D.W., Shatz, C.J., and Stryker, M.P. (1988). Modification of retinal ganglion cell axon morphology by prenatal infusion of tetrodotoxin. Nat. 1988 3366198 336, 468–471.

Sun, W., McConnell, E., Pare, J.-F., Xu, Q., Chen, M., Peng, W., Lovatt, D., Han, X., Smith, Y., and Nedergaard, M. (2013a). Glutamate -Dependent Neuroglial Calcium Signaling Differs Between Young and Adult Brain. Science (80-.). 339, 197–200.

Sun, W., McConnell, E., Pare, J.F., Xu, Q., Chen, M., Peng, W., Lovatt, D., Han, X., Smith, Y., and Nedergaard, M. (2013b). Glutamate-dependent neuroglial calcium signaling differs between young and adult brain. Science (80-.). 339, 197–200.

Tritsch, N.X., Yi, E., Gale, J.E., Glowatzki, E., and Bergles, D.E. (2007). The origin of spontaneous activity in the developing auditory system. Nature 450, 50–55.

Tritsch, N.X., Rodríguez-Contreras, A., Crins, T.T.H., Wang, H.C., Borst, J.G.G., and Bergles, D.E. (2010). Calcium action potentials in hair cells pattern auditory neuron activity before hearing onset. Nat. Neurosci. 13, 1050–1052.

Ventura, R., and Harris, K.M. (1999). Three-dimensional relationships between hippocampal synapses and astrocytes. J. Neurosci. 19, 6897–6906.

Voufo, C., Chen, A.Q., Smith, B.E., Yan, R., Feller, M.B., and Tiriac, A. (2023). Circuit mechanisms underlying embryonic retinal waves. Elife 12.

Wang, C., Wang, Y., and Yu, G. (2022). Efficient Global MOT under Minimum-Cost Circulation Framework. IEEE Trans. Pattern Anal. Mach. Intell. 44, 1888–1904.

Wang, Y., DelRosso, N. V., Vaidyanathan, T. V., Cahill, M.K., Reitman, M.E., Pittolo, S., Mi, X., Yu, G., and Poskanzer, K.E. (2019). Accurate quantification of astrocyte and neurotransmitter fluorescence dynamics for single-cell and population-level physiology. Nat. Neurosci. 1–9.

Wang, Y., Wang, C., Ranefall, P., Broussard, G.J., Wang, Y., Shi, G., Lyu, B., Wu, C.T., Wang, Y., Tian, L., et al. (2020). SynQuant: an automatic tool to quantify synapses from microscopy images. Bioinformatics 36, 1599–1606.

Winchenbach, J., Düking, T., Berghoff, S.A., Stumpf, S.K., Hülsmann, S., Nave, K.-A., and Saher, G. (2016). Inducible targeting of CNS astrocytes in Aldh1l1-CreERT2 BAC transgenic mice. F1000Research 5, 2934.

Wong, W.T., Sanes, J.R., and Wong, R.O.L. (1998). Developmentally Regulated Spontaneous Activity in the Embryonic Chick Retina. J. Neurosci. 18, 8839–8852.

Xu, H., Furman, M., Mineur, Y.S., Chen, H., King, S.L., Zenisek, D., Zhou, Z.J., Butts, D.A., Tian, N., Picciotto, M.R., et al. (2011). An Instructive Role for Patterned Spontaneous Retinal Activity in Mouse Visual Map Development. Neuron 70, 1115–1127.

Xu, H.P., Burbridge, T.J., Chen, M.G., Ge, X., Zhang, Y., Zhou, Z.J., and Crair, M.C. (2015). Spatial pattern of spontaneous retinal waves instructs retinotopic map refinement more than activity frequency. Dev. Neurobiol. 75, 621–640.

Xu, J., Zhu, Y., Contractor, A., and Heinemann, S.F. (2009). mGluR5 Has a Critical Role in Inhibitory Learning. J. Neurosci. 29.

