## Supplemental Figure 1 for "Conservation of neuron-astrocyte coordinated activity among sensory processing centers of the developing brain"

### Supplementary Figure 1

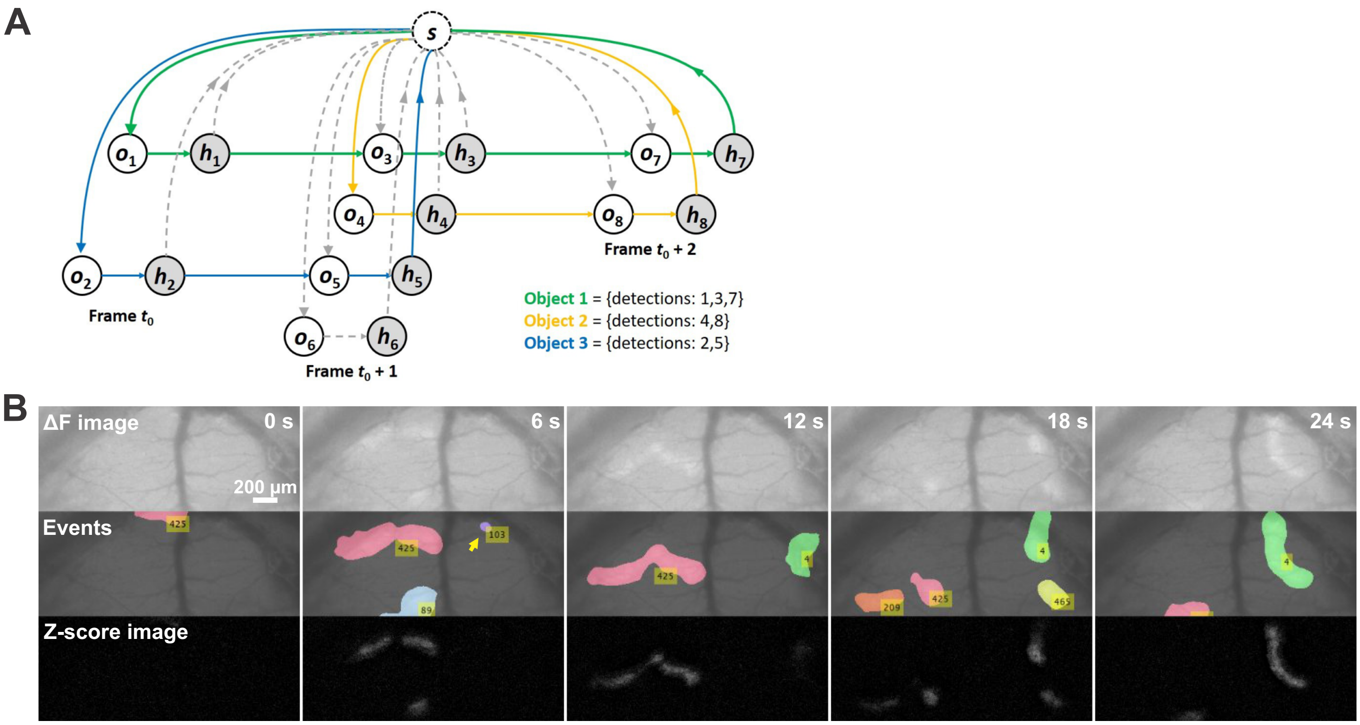


**Supplementary Figure 1. Tracking program for calcium waves in the superior colliculus**. **A**. Diagram of minimum-cost circulation graph. The edge from $o_i$ to $h_i$ is with negative weight representing the reward of each detection. The edges from $s$ to $o_i$ or from $o_i$ to $s$ are with positive weights representing the cost for flowing into/out one trajectory. The edge from $h_i$ to $o_j$ represents the cost flowing from $i_{th}$ detection to $j_{th}$ detection. **B**. Montage of detection of astrocyte calcium waves in the superior colliculus of a postnatal day 8 mouse. Top shows the ΔF image; Middle shows the events that were tracked in different colors (pseudocolored). Note that the yellow arrow points at an event that was manually removed by the user because it did not propagate. Bottom shows the Z-score image which was used for the detection algorithm.
